# The role of a non-native host plant in altering the seasonal dynamics of monarch development

**DOI:** 10.1101/2024.08.23.609406

**Authors:** James G. DuBose, Mackenzie Hoogshagen, Jacobus C. de Roode

## Abstract

The development and life history of many organisms have evolved to align with annual seasonal changes, but anthropogenic ecological changes have started to disrupt phenological dynamics. To understand the implications of these changes, tractable model systems are needed to identify the causes and consequences of phenological-seasonal asynchrony. Here, we investigate the potential of a non-native and tropical host plant whose phenology has been tuned to different seasonal dynamics to influence the seasonal developmental dynamics of the migratory monarch butterfly *Danaus plexippus*. Consistent with predictions, we found that the non-native host plant facilitated successful monarch development later into the fall and early winter, which is typically the season North American monarchs enter reproductive diapause and migrate to over-wintering locations. Furthermore, we found evidence that this success could partly be attributed to decreased thermal constraints on development experienced by monarchs reared on *A. curassavica*.

## Introduction

Seasonality represents one of the most pervasive sources of environmental variation that has shaped the life histories of organisms across the tree of life. Many organisms’ development is highly sensitive to environmental conditions, resulting in widespread evolution of developmental plasticity and phenological patterns that have been fine tuned to align with seasonal changes (Gilbert 2001; Visser et al. 2010; Moczek et al. 2011). However, anthropogenic environmental and ecological change has started interfering with major developmental and life history events, resulting in asynchrony between organisms’ phenologies and seasonal dynamics (Visser and Both 2005). Although many phenological shifts have been linked to changes in seasonal temperature patterns (Parmesan and Yohe 2003; Körner and Basler 2010), biotic factors can interact with seasonal temperature changes to influence the physiology of developing organisms, resulting in phenological shifts that are not predicted by temperature changes alone (Van Asch and Visser 2007; Gellesch et al. 2013; Rudolf and Singh 2013). Therefore, anthropogenic introduction of non-native species, whose phenology has been tuned to different seasonal dynamics, can play a significant role in generating asynchrony between organisms’ phenologies and seasonal dynamics (Renner and Zohner 2018). While this is a well described issue, tractable model systems for gaining a comprehensive understanding of the causes and consequences of phenological-seasonal asynchronies are limited (Renner and Zohner 2018).

The monarch butterfly *Danaus plexippus* has the potential to serve as an excellent model system for studying the causes and consequences of asynchrony between phenology and seasonal dynamics. Each year, millions of monarch butterflies undergo a seasonal migration that traverses most of North America (Reppert and de Roode 2018). This vast migration is facilitated by the development of several morphological and physiological traits, but most notably, reproductive diapause and the migratory behavior (Herman 1981). These traits are expressed in individuals that undergo metamorphosis at the end of breeding season (late summer-early fall), resulting in a seasonal migratory phenotype that is distinct form the parental generation (Alonso-Mejía et al. 1997; Schroeder et al. 2020). North American monarchs typically feed on the native milkweed species (*Asclepias spp*.) that senesce in the fall, which – combined with shortening day length and lower temperature – is thought to cue monarch migration by diminishing suitable habitat for reproduction and signaling the development of migrant traits during monarch metamorphosis (Goehring and Oberhauser 2002). However, the tropical milkweed *A. curassavica*, which does not seasonally senesce to the same extent or rate as native milkweeds (Woodson 1954; R.V. Batalden and K.S. Oberhauser 2015; Clement and Crawford 2020; James et al. 2021), has recently been introduced into North America and is widely commercially distributed.

Several studies have indicated a significant role of *A. curassavica* in promoting prolonged or year-round breeding in monarch populations (Satterfield et al. 2016, 2018; Clement and Crawford 2020). However, the physiological changes experienced by developing monarchs that underlie this phenomenon are less understood. Here, we compared the developmental success and thermal constraints on development of monarchs reared on *A. curassavica* and the native *A. incarnata* in the summer and fall. Overall, we found support for the hypothesis that *A. curassavica* facilitates successful monarch development later into the fall, as well as evidence that this success could be partly attributed to reduced thermal constraints on monarch development.

## Methods

### Study site and experimental design

To investigate the influence of season and host plant on monarch development, we reared monarchs on *A. curassavica* and *A. incarnata* and tracked their development during the summer and fall-winter seasons. We purchased the milkweed seeds used in this study form Joyful Butterfly (Blackstock, SC, USA) and propagated them in the greenhouse before re-planting at the outside study site (Atlanta, GA, USA, 33°44’55”N 84°23’15”W) approximately one month before the start of the study (May 2023). To obtain the larvae used in this study, we first collected wild-caught monarchs near St. Marks, Florida, U.S.A (30°09’33”N 84°12’26”W) to establish laboratory lines. Just prior to the start of the experiment in each season, we mated monarchs at the study site and transferred larvae directly to their treatment plants.

During the summer, we placed 2-3 larvae onto 29 *A. incarnata* and 29 *A. curassavica* plants, resulting in 86 *A. incarnata*-reared and 87 *A. incarnata*-reared monarchs. We placed the first larvae on their treatment plants starting on June 12, 2023, and continued to place larvae through June 18, 2023. A total of 38 larvae (19 from each plant species) and 36 pupae (17 from *A. curassavica* and 19 from *A. incarnata*) pupae were removed for another concurrent study, resulting in 49 *A. incarnata*-reared and 50 *A. curassavica*-reared monarchs that had the potential to reach adulthood. During the fall-winter, 2-3 larvae were placed onto 27 *A. curassavica* and 24 *A. incarnata* plants, resulting in 70 *A. incarnata*-reared and 80 *A. curassavica*-reared monarchs. We placed the first larvae on treatment plants October 4, 2023, and continued to place larvae through October 6, 2023. To ensure monarchs did not escape, we used a wooden steak and mesh net to construct a sealed enclosure around each plant. We placed HOBO Pendant data loggers in four of the enclosures to record the temperature every fifteen minutes throughout the duration of the experiment in each season.

### Data collection and analysis

Throughout the duration of the experiment in each season, we checked enclosures every day and recorded the dates of pupations and eclosions. Prior to the placement of larvae on treatment plants, we placed HOBO Pendant data loggers in four of the enclosures to record the temperature every fifteen minutes throughout the duration of the experiment in each season, which were averages for subsequent analysis. To gain insight into monarch development, we calculated growing degrees (GD) as: 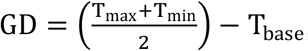, where T_base_ = 11.5°C (Zalucki 1982). We then calculated growing degree days 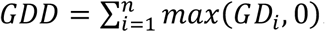. More generally, growing degree days give insight into how much heat an organism has accumulated over time, which is closely related to developmental dynamics in ectotherms.

We performed z-tests to assess differences in survival proportions on each plant and in each season. To test for difference in growing degree days, we used Wilcoxon rank-sum tests. We note that survival was too low in the *A. incarnata*-fall treatment for meaningful statistical comparisons, so this treatment was excluded from this analysis. All statistical analyses were performed in R (R Core Team 2022).

## Results

### The effects of host plant on seasonal developmental success

Because *A. curassavica* does not seasonally senesce at the same rate or extent as *A. incarnata*, we hypothesized that feeding on *A. curassavica* would allow for greater monarch survival later into the fall and winter. Consistent with this hypothesis, in the fall-winter season, the proportion of larvae that reached pupation when reared on *A. curassavica* (0.275) was greater than when reared on *A. incarnata* (0.1) (*X*^*2*^ = 6.252, p = 0.01241) (Figure 1A). Likewise, the proportion of pupae that eclosed when reared on *A. curassavica* (0.1375) was greater than when reared on *A. incarnata* (0.043) (*X*^*2*^ = 2.913, p = 0.08). However, the proportion of monarchs that reached pupation and eclosion in the fall-winter was still notably lower than during the summer (0.731 and 0.62, respectively) (pupation: *X*^*2*^ = 28.608, p = 8.86×10^−8^; eclosion: *X*^*2*^ = 30.585, p = 3.195×10^−8^), where the proportions were higher and comparable between both host plant species (Figure 1).

**Figure 1.**
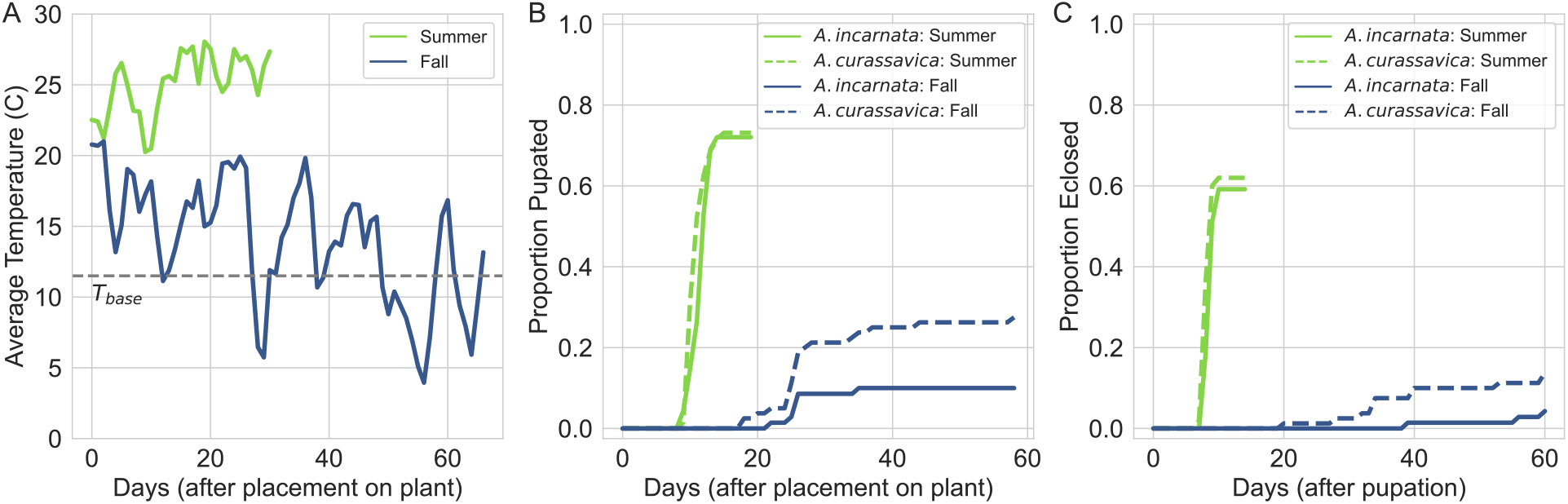
*A. curassavica* facilitates monarch development later into the fall than *A. incarnata*. A) Average daily temperature as a function of time for summer and fall conditions. The dashed line shows the base temperature required for monarch development. B and C) Development curves as a function of time for larval development into pupae and pupal development into adults, respectively. In B, the x-axis indicates the number of days between the placement of larvae on their treatment plants and pupation. In C, the x-axis indicated the number of days between pupation and eclosion. In B and C, proportions on the y-axis indicate the proportion of total monarchs reared in each season that reached pupation and eclosion, respectively.

### The effects of plant host on thermal constraints of development

We then asked if plant host influenced thermal constraints on development (Figure 2A), and if this varied by season and life stage. We analyzed thermal constraints on larval and pupal development separately because they have distinct biological functions. The larval stage is important for accumulating energy and storing fat, while the pupal stage is responsible for phenotypic transitioning and maturation into reproductive adults. We found that monarchs reared on *A. curassavica* during the summer completed development with less thermal input than those reared on *A. incarnata* in the summer (W = 235.5, p = 0.0008134) (Figure 2A); this effect was apparent in larval (W = 869, p = 0.005078) (Figure 2B) and pupal (W = 306.5, p = 0.02187) development (Figure 2C). During the fall, monarchs reared on *A. curassavica* were able to complete development despite having significantly less thermal input than those reared on *A. curassavica* during the summer (W = 8, p = 3.723 x 10^−5^) (Figure 2A). Although we cannot make comparisons to monarchs reared on *A. incarnata* due to low survivorship to adulthood, these findings suggest a role of *A. curassavica* in reducing thermal constraints on completing metamorphosis.

**Figure 2.**
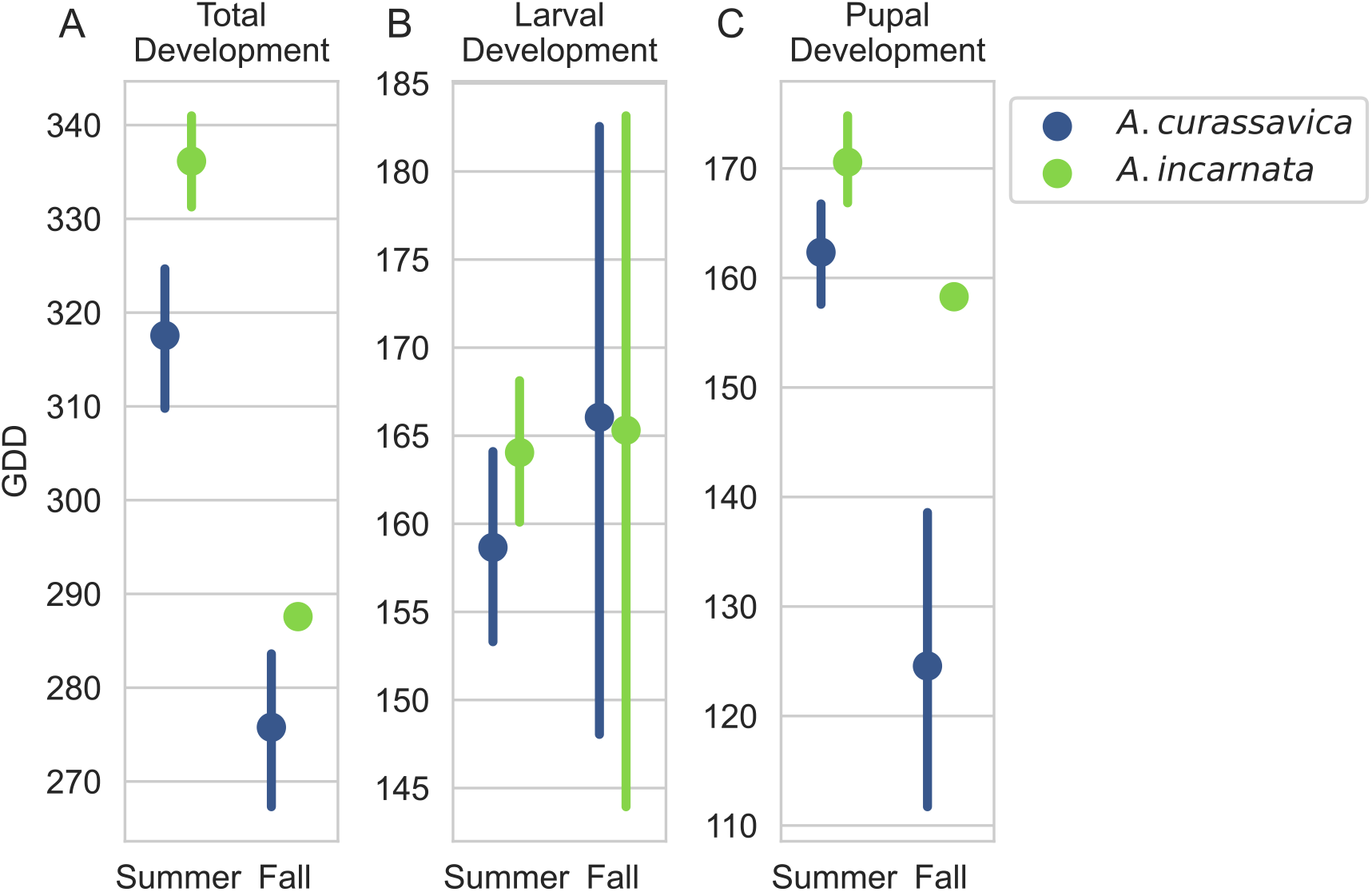
Monarchs reared on *A. curassavica* complete development with less thermal input. Point plot representing the accumulated growing degrees (GDD) during A) total development, B) larval development, and C) pupal development. Points represent the mean GDD, and lines represent the 95% confidence interval.

## Discussion

Understanding the causes and consequences of seasonal (a)synchronies in phenology is essential for predicting how populations will respond to anthropogenic environmental and ecological change, but the physiological/developmental bases of said asynchronies are less understood (Van Asch and Visser 2007). Here, we investigated the consequences of *A. curassavica*, a non-native host plant that does not seasonally senesce as readily as native hosts (Woodson 1954; R.V. Batalden and K.S. Oberhauser 2015; Clement and Crawford 2020; James et al. 2021), on the seasonal dynamics of monarch developmental. We found that feeding on *A. curassavica* facilitated monarch development later into the fall/winter, as well as evidence that this success could be partly facilitated by decreased thermal constraints on development experienced by monarchs that were reared on *A. curassavica*. Furthermore, monarchs reared on *A. curassavica* were able to complete their metamorphosis despite prolonged periods where the average temperature dropped below their development zero (T_base_, Figure 2A).

Our finding that *A. curassavica* facilitated monarch development later into the fall/winter is consistent with natural observations that increased monarch fall/winter breeding is associated with prolonged availability and density of *A. curassavica* as a food source (Howard et al. 2010; R.V. Batalden and K.S. Oberhauser 2015; Kendrick and McCord 2023; Steele et al. 2023). Furthermore, previous experimental work has found that feeding on *A. curassavica* as larvae is linked to the development of reproductive traits as adults, and that exposure to *A. curassavica* can break reproductive diapause, lending further evidence that *A. curassavica* has a role in promoting fall/winter breeding (Satterfield et al. 2018; Majewska and Altizer 2019). Our finding that feeding on *A. curassavica* was associated with reduced thermal constraints on metamorphosis expands on this previous work by describing a potential physiological basis for successful monarch development in colder conditions. However, it is important to note that this study is limited in that we were not able to directly compare monarch thermal accumulation between *A. curassavica* and *A. incarnata* in the fall/winter due to low survivorship on *A. incarnata*. Future work that focuses on establishing causality and mechanistically validating these findings, as well as describing the continuous relationship between the degree of seasonal plant senescence and monarch developmental dynamics, will be useful in assessing this hypothesis.

More generally, phenological asynchronies re-shape a variety of ecological interactions (Van Asch and Visser 2007; Schwartzberg et al. 2014; Uelmen et al. 2016). In the context of monarchs, this phenomenon is best described in their interaction with a protozoan parasite *Ophryocystis elektroscirrha*. Sedentary monarch sub-populations, which do not undergo seasonal migration, have been found to have higher parasite prevalence (Satterfield et al. 2015, 2016). This can in turn increase parasite infection in migratory monarchs when they interact with sedentary monarchs (Satterfield et al. 2018). Overall, this illustrates that predicting the consequences of phenological asynchronies requires integrating understanding of its implications for population and ecological dynamics.

## Data availability

All data collected and code written for this project can accessed at https://github.com/gabe-dubose/w3mp.

## Acknowledgements

We thank Erik Edwards for initial growing the plants that were transplanted to the field site, as well as the members of the de Roode lab for helping manage monarch mating. This work was supported by National Science Foundation grant IOS-1922720 to J.C.dR.

## Conflict of interest

The authors declare no conflict of interest.

## Notes

https://github.com/gabe-dubose/w3mp

### Competing Interest Statement

The authors have declared no competing interest.

## References

Alonso-Mejía, A., E. Rendon-Salinas, E. Montesinos-Patiño, and L. P. Brower. 1997. USE OF LIPID RESERVES BY MONARCH BUTTERFLIES OVERWINTERING IN MEXICO: IMPLICATIONS FOR CONSERVATION. Ecological Applications 7:934–947.

Clement, K., and P. Crawford. 2020. FALL AVAILABLE TROPICAL MILKWEED (ASCLEPIAS CURASSAVICA L.) MAY BE A POPULATION SINK FOR THE MONARCH BUTTERFLY. onpr 20.

Gellesch, E., R. Hein, A. Jaeschke, C. Beierkuhnlein, and A. Jentsch. 2013. Biotic Interactions in the Face of Climate Change. Pp. 321–349 in U. Lüttge, W. Beyschlag, D. Francis, and J. Cushman, eds. Progress in Botany. Springer Berlin Heidelberg, Berlin, Heidelberg.

Gilbert, S. F. 2001. Ecological Developmental Biology: Developmental Biology Meets the Real World. Developmental Biology 233:1–12.

Goehring, L., and K. S. Oberhauser. 2002. Effects of photoperiod, temperature, and host plant age on induction of reproductive diapause and development time in Danaus plexippus. Ecological Entomology 27:674–685.

Herman, W. S. 1981. Studies on the adult reproductive diapause of the monarch butterfly, Danaus plexippus. The Biological Bulletin 160:89–106.

Howard, E., H. Aschen, and A. K. Davis. 2010. Citizen Science Observations of Monarch Butterfly Overwintering in the Southern United States. Psyche: A Journal of Entomology 2010:1–6.

James, D. G., M. C. Schaefer, K. Krimmer Easton, and A. Carl. 2021. First Population Study on Winter Breeding Monarch Butterflies, Danaus plexippus (Lepidoptera: Nymphalidae) in the Urban South Bay of San Francisco, California. Insects 12:946.

Kendrick, M. R., and J. W. McCord. 2023. Overwintering and breeding patterns of monarch butterflies (Danaus plexippus) in coastal plain habitats of the southeastern USA. Sci Rep 13:10438.

Körner, C., and D. Basler. 2010. Phenology Under Global Warming. Science 327:1461–1462.

Majewska, A. A., and S. Altizer. 2019. Exposure to Non-Native Tropical Milkweed Promotes Reproductive Development in Migratory Monarch Butterflies. Insects 10:253.

Moczek, A. P., S. Sultan, S. Foster, C. Ledón-Rettig, I. Dworkin, H. F. Nijhout, E. Abouheif, and D. W. Pfennig. 2011. The role of developmental plasticity in evolutionary innovation. Proc. R. Soc. B. 278:2705–2713.

Parmesan, C., and G. Yohe. 2003. A globally coherent fingerprint of climate change impacts across natural systems. Nature 421:37–42.

R Core Team. 2022. R: A Language and Environment for Statistical Computing. R Foundation for Statistical Computing, Vienna, Austria.

Renner, S. S., and C. M. Zohner. 2018. Climate Change and Phenological Mismatch in Trophic Interactions Among Plants, Insects, and Vertebrates. Annu. Rev. Ecol. Evol. Syst. 49:165– 182.

Reppert, S. M., and J. C. de Roode. 2018. Demystifying Monarch Butterfly Migration. Current Biology 28:R1009–R1022.

Rudolf, V. H. W., and M. Singh. 2013. Disentangling climate change effects on species interactions: effects of temperature, phenological shifts, and body size. Oecologia 173:1043–1052.

R.V. Batalden and K.S. Oberhauser. 2015. Potential changes in eastern North American monarch migration in response to an introduced milkweed, Asclepias curassavica. P. in Monarchs in a Changing World. Cornell University Press, Ithaca NY.

Satterfield, D. A., J. C. Maerz, and S. Altizer. 2015. Loss of migratory behaviour increases infection risk for a butterfly host. Proc. R. Soc. B. 282:20141734.

Satterfield, D. A., J. C. Maerz, M. D. Hunter, D. T. T. Flockhart, K. A. Hobson, D. R. Norris, H. Streit, J. C. De Roode, and S. Altizer. 2018. Migratory monarchs that encounter resident monarchs show life-history differences and higher rates of parasite infection. Ecology Letters 21:1670–1680.

Satterfield, D. A., F. X. Villablanca, J. C. Maerz, and S. Altizer. 2016. Migratory monarchs wintering in California experience low infection risk compared to monarchs breeding year-round on non-native milkweed. Integr. Comp. Biol. 56:343–352.

Schroeder, H., A. Majewska, and S. Altizer. 2020. Monarch butterflies reared under autumn-like conditions have more efficient flight and lower post-flight metabolism. Ecological Entomology 45:562–572.

Schwartzberg, E. G., M. A. Jamieson, K. F. Raffa, P. B. Reich, R. A. Montgomery, and R. L. Lindroth. 2014. Simulated climate warming alters phenological synchrony between an outbreak insect herbivore and host trees. Oecologia 175:1041–1049.

Steele, C., I. G. Ragonese, and A. A. Majewska. 2023. Extent and impacts of winter breeding in the North American monarch butterfly. Current Opinion in Insect Science 59:101077.

Uelmen, J. A., R. L. Lindroth, P. C. Tobin, P. B. Reich, E. G. Schwartzberg, and K. F. Raffa. 2016. Effects of winter temperatures, spring degree-day accumulation, and insect population source on phenological synchrony between forest tent caterpillar and host trees. Forest Ecology and Management 362:241–250.

Van Asch, M., and M. E. Visser. 2007. Phenology of Forest Caterpillars and Their Host Trees: The Importance of Synchrony. Annu. Rev. Entomol. 52:37–55.

Visser, M. E., and C. Both. 2005. Shifts in phenology due to global climate change: the need for a yardstick. Proc. R. Soc. B. 272:2561–2569.

Visser, M. E., S. P. Caro, K. Van Oers, S. V. Schaper, and B. Helm. 2010. Phenology, seasonal timing and circannual rhythms: towards a unified framework. Phil. Trans. R. Soc. B 365:3113–3127.

Woodson, R. E. 1954. The North American Species of Asclepias L. Annals of the Missouri Botanical Garden 41:1.

Zalucki, M. P. 1982. Temperature and Rate of Development in Danaus plexippus L. and D. chrysippus L. (Lepidoptera: Nymphalidae). Australian Journal of Entomology 21:241–246.

